# Computational modeling of social face perception in humans: Leveraging the active appearance model

**DOI:** 10.1101/360776

**Authors:** Jinyan Guan, Chaitanya K. Ryali, Angela J. Yu

**Affiliations:** Department of Cognitive Science, University of California San Diego, 9500 Gilman Drive La Jolla, CA 92093; Department of Computer Science and Engineering, University of California San Diego, 9500 Gilman Drive La Jolla, CA 92093

## Abstract

Face processing plays a central role in human social life. Humans readily infer social traits (e.g. attractiveness and trustworthiness) from a stranger’s face. Previous attempts to characterize the facial (physiognomic) features underlying social processing have lacked either systematicity or interpretability. Here, we utilize a statistical framework to tackle this problem, by learning a vector space to represent faces, and a linear mapping from this face space into human social trait judgments. Specifically, we obtain a face space by training the Active Appearance Model on large datasets of face images. Based on human evaluations of numerous social traits on these images, we then use regression to find linear combinations of facial features (what we call Linear Trait Axis, or LTA) that best predict human social judgments. Our model achieves state-of-the-art performance in overall predictive accuracy – comparable to the best convolutional neural network and better than human prediction of other human ratings. To interpret the LTAs, we regress them against a large repertoire of geometric features. To understand the relationship between the facial features that underlie different social, emotional, and demographic traits, we present a novel “dual space analysis” that characterizes the geometric relationship among LTA vectors. It shows that facial features important for social trait perception are largely distinct from those underlying demographic and emotion perception, contrary to previous suggestions that social trait perception is driven by over-generalization of relatively primitive demographic and emotion perception processes. In addition, we present a novel correlation decomposition analysis that quantifies how correlations in trait judgments (e.g. between attractiveness and babyfacedness) independently arise from (1) shared facial features among traits, and (2) correlation in the distribution of facial features in the human population.

## 1 Introduction

Humans readily infer social traits, such as attractiveness and trustworthiness, by glancing at a stranger’s face [1, 2]. While the veracity of these judgments is an area of active research [3, 4], they certainly exert a powerful influence on real-life decisions [5], e.g. choosing a mate, assessing witness testimony, interviewing a candidate, deciding whom to befriend in a crowd. Understanding which facial (physiognomic) features drive the perception of different personality traits is important for advancing basic neuroscience and psychology, as well as improving engineered systems that need to anticipate or generate faces with desired human social perception.

Several lines of previous research have attempted to characterize the facial (physiognomic) features underlying social trait judgments, but they typically lacked either systematicity or interpretability. One classical approach has been motivated by the notion that social impressions of faces are over-generalizations of relatively primitive face processing elements. For example, babyfacedness [6, 7, 8] and facial width-to-height ratio [9, 10, 11] are thought to be important for the perception of kindness and aggressiveness, respectively. Previous work, by morphing faces to different emotion expressions, has suggested that emotional aspects of the face can drive social impressions [12]. While hypothesis driven, this approach is ad hoc and lacks systematicity. It cannot, for example, elucidate what facial features besides the hypothesized one(s) may drive the perception of a social trait. Another line of work applies principal component analysis (PCA) to simultaneous judgment of multiple traits, and presumes that the traits most closely aligned with the principal axes to be “fundamental” traits. This approach has identified trustworthiness (also known as “valence”) and dominance as the two primary traits [13, 14], and “youthful-attractiveness” as a possible third dimension [15, 16]. However, as this approach only analyzes human ratings and not face images, it does not clarify which facial features actually drive the perception of the “fundamental” traits, nor what factors besides the “fundamental” traits may contribute to the perception of a specific trait. A third approach, using a computational model that attempts to parameterize all faces [17, 18], can generate synthetic faces that vary in social impression ratings [19, 20, 21, 22]. However, this approach is challenging for interpretation, as it requires ad hoc visual inspection of the synthetic images to generate verbal description of the features that matter for each trait. Another weakness is that it is unclear whether these models have a representation rich enough to capture all ethologically relevant variability in facial features. Both of these problems are exacerbated by the common use of a popular but opaque commercial software, FaceGen (Singular Inversions, https://facegen.com/), in the literature.

In this work, we build on this third line of work, by learning a vector space representation of faces using the Active Appearance Model (AAM) [23, 24], and a linear mapping between this face space and human social trait judgments. In section 2, we motivate, describe and validate our model. In section 3, we present three novel methods for analyzing and interpreting the facial features underlying social trait perception, as well as how the facial features underlying different traits are related to each other. In section 4, we discuss our contributions, broader impact on the field, shortcomings, and future directions of research.

## 2 Face model training and evaluation

There are two distinct components to our model of human social perception of faces. The first is *unsupervised learning* of a face representation that ideally encompasses all natural variations among face images. The second is *supervised learning* of a mapping between this generic face space and human ratings for different social traits (e.g. attractiveness, trustworthiness). In the following, we describe these two components in detail, followed by experimental validation of the model.

### 2.1 Unsupervised learning of face representation

Several properties of a vector space representation of faces [25] are particularly desirable: (1) we want a sufficiently complex representation, such that each real face image maps to a point (ideally unique) in this space; (2) we want it to be a generative model, such that each point generates a realistic face image; (3) we want the dimensionality of the space to be small enough to enable useful analyses; (4) we want the representation to be transparent enough so that we can gain some insight into the facial (physiognomic) features that underlie human perception of different traits; (5) an added bonus is if the representation has some neural relevance, such that it allows us not only to model human *behavior*, but potentially also the neural basis underlying social face representation and processing.

The above desiderata led us to AAM, a well-established machine vision technique that does a good job of reconstructing images (property 1), generates realistic synthetic faces (property 2), and has a latent representation of only a few dozen features (property 3) [23, 26, 24]. Additionally, it has been shown that AAM features are well encoded by face processing neurons in the monkey brain, and that this neuronal representation is rich enough to discriminate different individuals [27]. Finally, AAM also has a fairly transparent representation (property 4) as follows. Each face image defines a *shape vector*, which is just the (x,y) coordinates of some consistently defined landmarks across all faces – in our case, we use the free software Face++ (https://www.faceplusplus.com), which labels 83 landmarks (e.g. contour points of the mouth, nose, eyes). Each face image also defines a *texture vector*, which is just the grayscale pixel values of a warped version of the image after aligning the landmark locations to the average landmark locations across the data set. The dataset we use consists of 2222 US adult face images that are collected from Google Images (D-1) [28], and 597 face images with neutral facial expression taken in the laboratory (D-2) [29]. We learned an AAM that contains a set of basis that describe the joint variations in shape and texture and principal components (PC) representing the variation of the shape and texture of a face from the average face.

This learned AAM allows us to *generate* any synthetic face in the face space: we take the mean face image, obtained by the average shape and the pixel-wise averaging of the landmark-aligned (shape-free) versions of all the training faces, and then add to it shape and feature distortions implied by each PC with a weight given by the PC coefficient.

### 2.2 Supervised learning of social trait judgment

For simplicity, we use multiple linear regression to find the best linear mapping from the face space (spanned by the orthogonal PCs) to average human ratings for each trait. Each face image in D-1 [28] comes with 15 individual Amazon M-Turk ratings (1-9) along 20 social traits and their antonyms (see Figure 3E;F); to deal with subjective differences in social perception or interpretation of the scale, we standardize ratings across images within each rater for each trait. In addition, D-1 face images come with modal binary ratings of gender, modal categorical ratings of race (which we turn into one-hot encoding), as well as average real-valued ratings of age. Each face image in D-2 [29] comes with average ratings (mean: 46 ratings/image) on 9 social traits and 6 emotional expressions (see section 3). For each trait, we linearly regress all the ratings against the first 70 PCs in the learned AAM(see section 2.1?), a dimensionality that minimizes the average cross-validation error (RMSE of ratings) across all D-1 social traits. We denote the normalized version of the regression coefficients (not including the constant term) the Linear Trait Axis (LTA, see Figure 3A): it is also the best linear combination of facial features for accounting for variations in a trait, and it also specifies the optimal linear gradient for increasing/decreasing the perception of a trait.

More formally, a subject’s rating on a specific social traits is given by *y* = x^T^ *β* + *β*_0_ + *ϵ*, where x *ϵ* ℝ^70^, where *ϵ* is a Gaussian distributed random variable with zero mean and some variance that are shared by all faces. We define the linear trait axis (LTA) as a normalized vector of the regression coefficients: 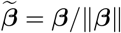. We can use ordinary least squares to estimate the coefficients *β* for each trait. Then the predicted rating of a subject is given by *ŷ* = x^T^ *β* + *β*_0_. Using the LTAs, we can synthesize faces that exhibit certain trait perception by changing a base face along the corresponding LTA direction.

### 2.3 Model evaluation

To visualize the most prominent features of AAM, we generate synthetic faces along the first three PCs (Figure 1A): PC 1 appears to capture both extrinsic environmental variables like illumination and intrinsic face variables like complexion. PC 2 appears to mostly capture rotation along the horizontal axis. PC 3 appears to capture holistic properties about the shape of the face. Since we can embed all faces distinctly into the AAM feature space, we can compute the sample statistics of all the training data, and thus generate faces that are two standard deviations away from the mean face in either direction for each PC. We therefore have a *statistical* model of the face space.

**Figure 1:**
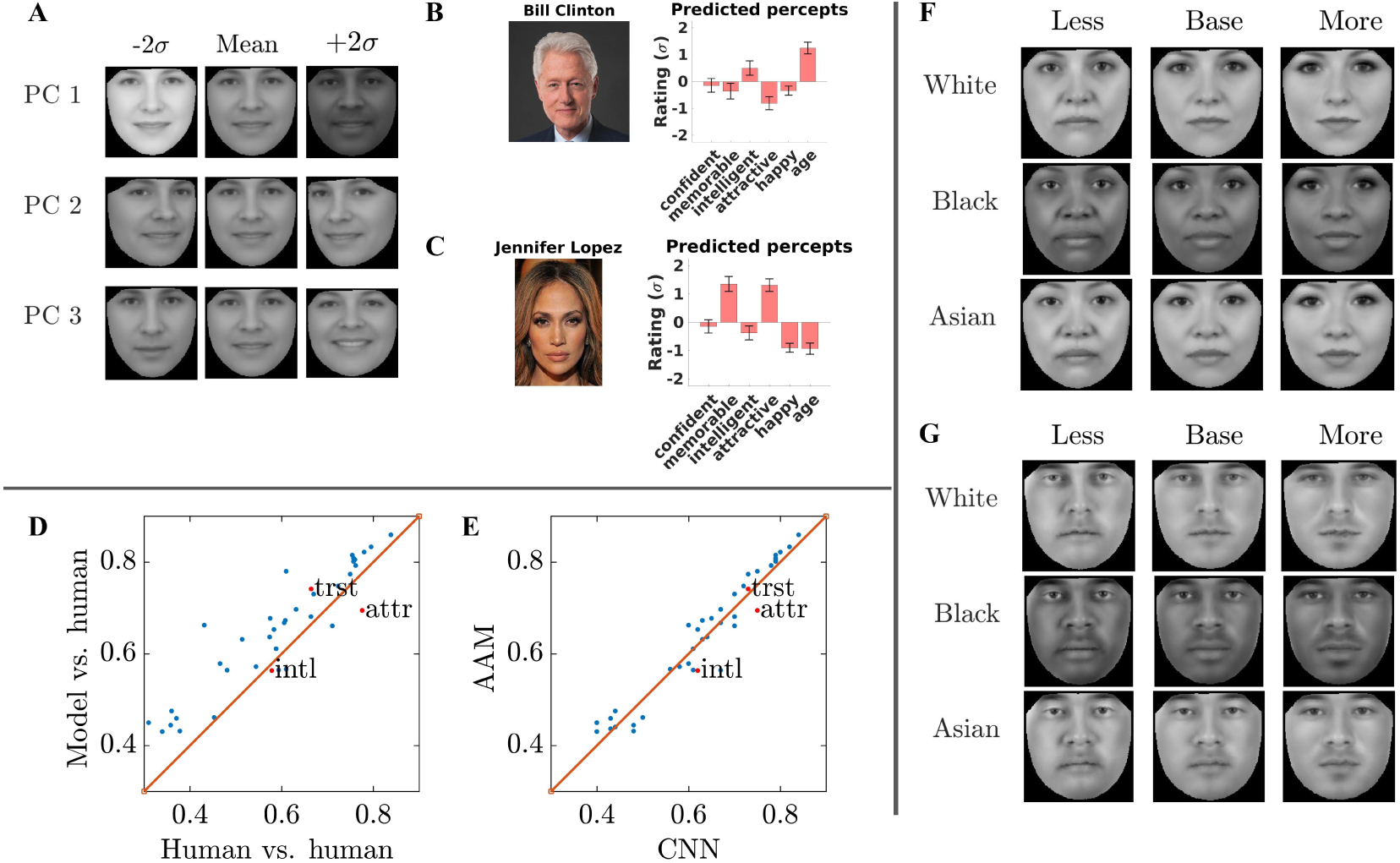
Modeling social trait perception with AAM. (A) Synthetic faces generated along the first three AAM PCs: middle - mean face, left and right: two standard deviations away from the mean face. (B) Predicted ratings (in units of training data std in D-1) on various traits for a Bill Clinton image. (C) Simlar to (B) for a Jennifer Lopez image. (D) Correlation between model-predicted social traits and average human ratings on a novel face (y-axis) vs. correlation between the average rating of one group of (7-8) human raters an another group of (7-8) human raters. (E) Same as (D) for the y-axis, and correlation between CNN-predictions and average human ratings (x-axis). (F-G) Synthetic faces along attractive LTA (right: more attractive, left: less attractive) for different base faces.

Using our model, we can predict social trait perception for any novel face. Figure 1B;C shows two such examples (images of Bill Clinton and Jennifer Lopez). Because we have a statistical model of faces based on the large training set, we can compute not only a mean prediction in units of standard deviation relative to the population, but also the 95% confidence interval (error bars) using the learned regression model. For example, we can say that this Bill Clinton face image is significantly above population mean (t-test, *α* = 0.05 significance level) in perceived “intelligence” and “age”, and below population mean in perceived “memorability”, “attractiveness”, and “happiness”, and that perceived “age” is about one standard deviation above population mean.

To validate the prediction performance of our model, we compare how well our model predicts human ratings (Figure 1D (y-axis)) on novel images to how well humans can predict other individuals’ ratings on the same image (x-axis) – we find that our model actually achieves better performance than humans. We suspect that this is because there are large individual variations in human social perception, and therefore ratings from one small group of people (7-8) may not be great for predicting ratings from another small group of people (7-8). Our model can do better because it can embed the novel face into a vector space, and leverage what it has learned about the social perception based on other faces, in particular what facial features matter and how, to make a good prediction. We also find that the trait prediction performance of our model (Figure 1E (y-axis)) is comparable to the state-of-the-art convolutional neural network (Figure 1E (x-axis)) [30].

Finally, we can use our model to take any “base” face, and apply the learned LTA for a trait to modify that base face in order to increase or decrease the perception of that trait. For example, Figure 1F shows how we can make female white, black, and Asian faces look more (right) or less (left) attractive; Figure 1G shows the same for male base faces. Note that these bases are also synthetically generated, by moving along the gender LTA. This is a visual illustration of how our model can successfully model social trait perception. We can of course apply the same manipulation to all the other traits we modeled (not shown due to space limitation).

## 3 LTA Analysis

Now that we have established by various means that the AAM-based linear models can successfully capture variations in human perception of social traits, we attempt to interpret the facial features that actually (linearly) drive the perception of each trait by analyzing the LTA’s. We present three analyses in the following three subsections.

### 3.1 Geometric feature analysis

The first method we introduce is to interpret the LTA in terms of quantitative and objective geometric features. The D-2 face images come with human-measured geometric features [29]. To illustrate the method, we regress the two example LTAs (attractiveness and babyfacedness) against the geometric features (Figure 2). We do so *sequentially*, in order to deal with correlations among the geometric features. It means that we first regress model-predicted attractiveness rating against the first feature, then take the residual and regress against the second, etc. The features are ordered so as to have decreasing absolute correlation coefficients with the model-predicted attractiveness rating. Thus, we first attribute any shared variance to a feature that explains more of the variation in trait rating. The features shown are only those that have significance regression coefficients in the sequential process.

**Figure 2:**
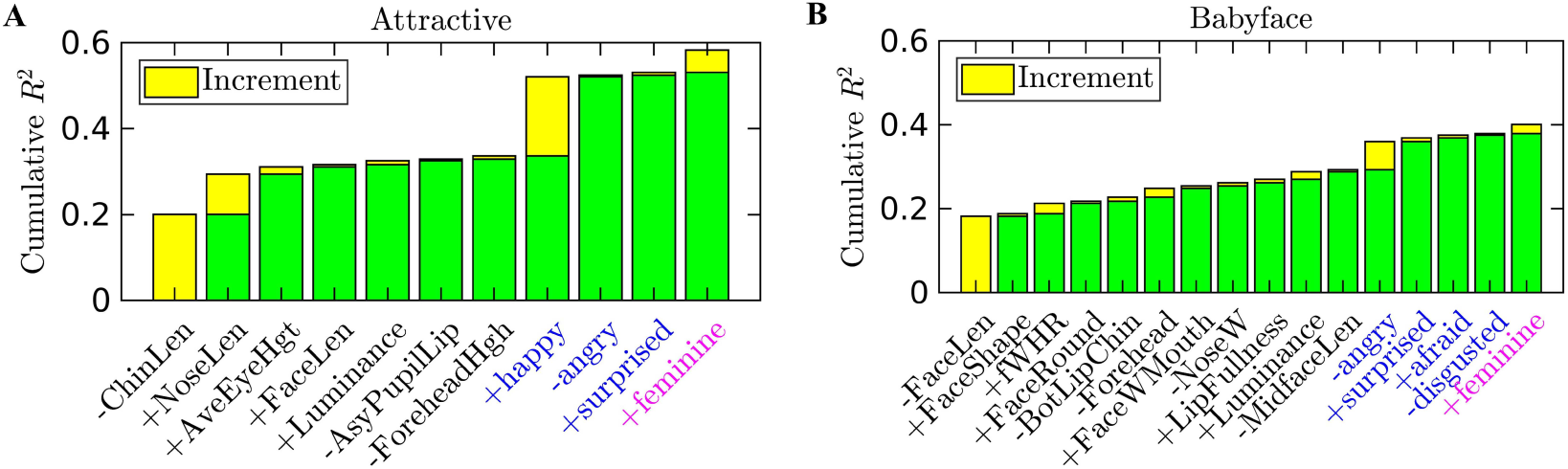
Cumulative *R*^2^ from sequential regression of attractiveness (A) and babyfacedness (B) against geometric features, LTA-predicted emotional traits, and LTA-predicted femininity perception. The geometric features are ordered in decreasing correlation coefficients with the model-predicted attractiveness. The ordering within emotional traits are done the same way.

We see that the major contributors to attractiveness LTA are decrease in chin length, increase in nose length, and there is a minor contribution by eye height (distance between upper and lower eyelids), and even smaller contribution by the other features (Figure 2A). Since the total *R*^2^ due to the geometric was not very high, we wondered if there were elements of attractiveness captured by emotional expressions (happiness, anger, surprise, disgust, fear, anger) and demographic traits (perceived femininity) that were not already present in the geometric features. Following the same sequential regression procedure, we see that a happy demeanor makes a sizable positive contribution to attractiveness, while negative emotional expressions (anger, surprise) make small but signficant negative contributions. Finally, looking more feminine appears to make a face more attractive, which is a little puzzling; but as we will see later on, this positive contribution is driven solely by the perception of female attractiveness, while male attractiveness does not depend on perception of femininity (at least in our linear model). Based on a similar analysis, we see that decreases in face length is the major geometric feature contributor to perceived babyfacedness (Figure 2B). In addition, looking less angry and more feminine both contribute to perceived babyfacedness.

Overall, the measured repertoire of geometric features [29] explains a significant but not overwhelming amount of the model-predicted trait variations (total *R*^2^ of geometric features for attractiveness: 0.35, babyfacedness: 0.31, femininity: 0.45, masculinity: 0.49, trustworthiness: 0.25). The significant residual correlation with some emotional expressions and femininity is interesting but still problematic for interpreting the LTAs, since we cannot explicitly characterize what facial features contribute to these emotional and demographic traits.

### 3.2 Dual Space Analysis

Given that the LTAs are themselves vectors that live in ℝ^70^ (Figure 3B), we can perform some geometric analyses of this dual space of LTAs to understand how the perception of different social traits relates to one another.

**Figure 3:**
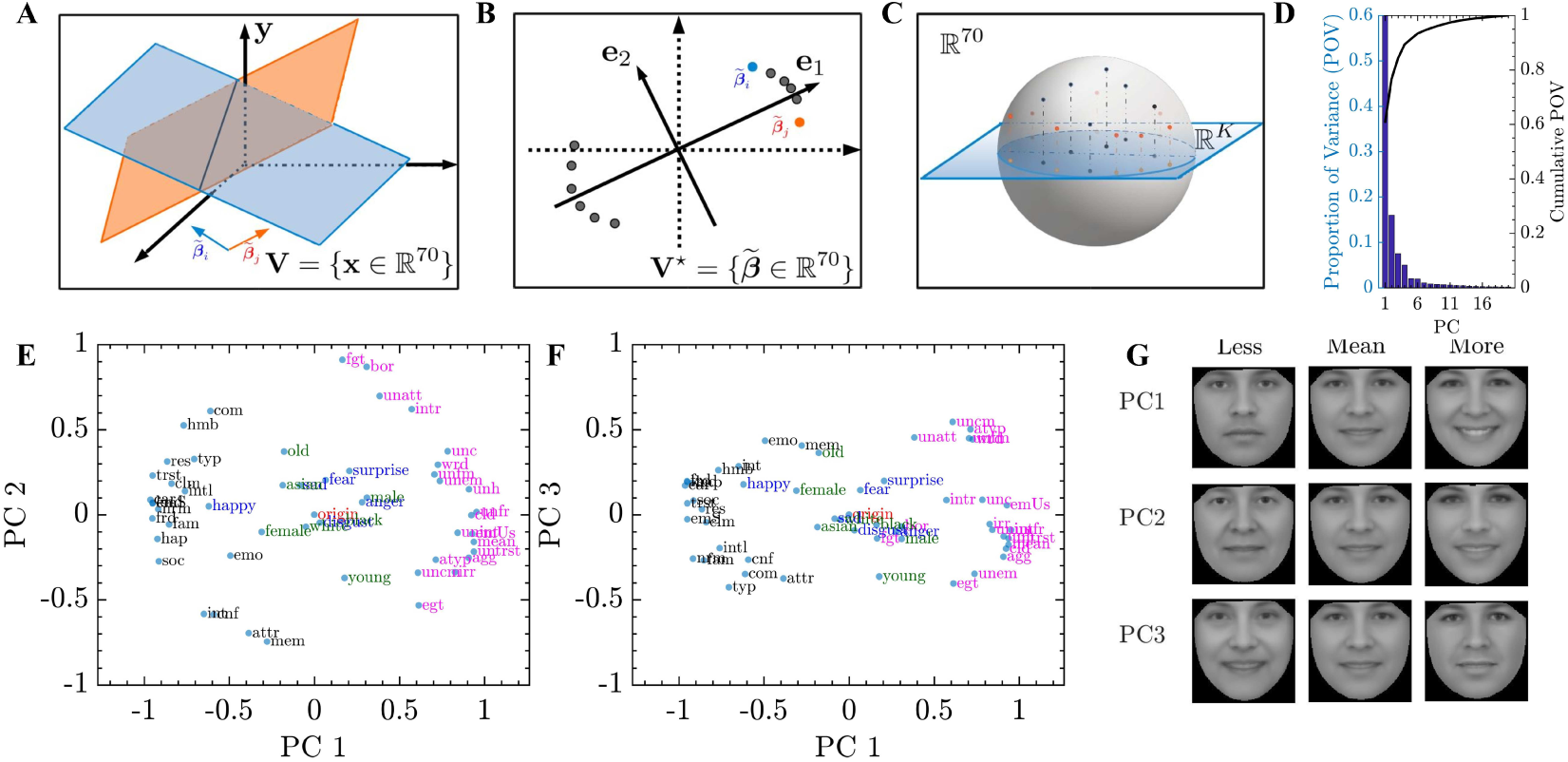
Dual space analysis: (A) Schematic illustration of linear maps for two traits, and their corresponding LTAs. (B) Schematic illustration of PCA of social LTAs in the dual space. (C) Schematic illustration of projecting LTAs down to a small subspace. (D) Incremental and cumulative proportion of variance explained by the PCs of social LTAs. (E) Projection of social LTAs (black: positive, magenta: negative), emotional LTAs (blue) and demographic LTAs (green) to the subspace spanned by the first two PCs of a PCA performed on the social LTAs. (F) Same as (F) but PC 1 and PC 3. (G) Synthetic faces along the first three PCs of the social LTAs.

Firstly, since social perception of traits exhibits high correlation among the traits [28], we suspect that the LTAs actually live in a much smaller dimensional subspace of ℝ^70^, i.e. there is strong overlap in the facial features that contribute to the perception of different social traits. We thus apply PCA to the 40 social traits from D-1 [28] (Figure 3B schematically illustrates this procedure). If the social LTAs mostly lie in a small subspace, then we expect the percent of variance (POV) explained by the PCs would rapidly decrease. This is indeed the case (Figure 3D): the first 3 PCs explain 83.46% of the variance, the first 5 PCs explain 95.1% of the variance (Figure 1F). We note that these LTA PCs are also just ℝ^70^ vectors corresponding to “principal” face feature axes that explain most of the (model-predicted) social traits. We can therefore generate synthetic faces to visualize these social “principal” LTA axes (Figure 3G). Unlike the feature axes in the original AAM space (Figure 1A), these principal LTA axes do not encode changes in nuisance image features (e.g. illumination, head orientation). As one would expect, human observers mainly care about subtle variations in intrinsic facial features when making social judgments. Another interesting implication of this analysis is that the number of facial features (orthogonal axes in the AAM face space) needed to explain human perception of a large variety of social traits [28] is actually far smaller than 70. Based on the POV analysis, a handful is already quite sufficient!

The next thing we analyzed is how social trait perception relates to demographic traits and emotional expressions. Social psychologists have suggested that social trait perception may be driven by demographic perception (in particular gender, age, and race) or are over-generalizations of the emotion perception system [31]. We can project the LTAs for the various social, demographic, and emotional traits down to the 3-dimensional subspace spanned by the first three PCs of the social LTAs. As shown in Figure 3C, LTAs that live close to this 3-dimensional subspace will have a projection close to the unit sphere (recall all LTAs have unit length), while those that are far away will have a projection close to the origin. Figure 3E;F show that PC 1 separates all the positive social traits from the negative ones, while PC 2 seems to account mostly for memorable/attractive versus forgettable/boring, and PC 3 is a harder to interpret dimension that has perhaps something to do with how emotional a face appears. In contrast to the social LTAs, which mostly lie close to the unit sphere (indicating good representation, as they well should), emotional and demographic traits (including gender) lie rather close to the origin, indicating they rely on facial features rather distinct from those driving social trait perception. The two exceptions are happiness LTA, which points almost directly along PC1 toward the center of the positive social LTAs and has magnitude close to 1, and young/old, which has relatively good representation in this subspace (indicating it relies many of the same facial features as social traits) but interestingly does not point in the same direction as any of the measured social traits.

### 3.3 Correlation decomposition

Finally, we use the LTAs to get a deeper understanding of how correlations in the perception of multiple traits arise. For example, the classical “attractiveness halo” [31] refers to the phenomenon that people who are judged attractive are also attributed with all kinds of positive personality traits, such as friendliness, trustworthiness, kindness, extroversion, etc. It has been implicitly assumed, rather universally, that such correlations between two traits arise because of common facial features that are important for both traits. However, our LTA representation raises the possibility of another, distinct source of perceptual correlation that has nothing to do with shared facial features, and that is *data correlation*.

It is obvious how common facial features can induce judgment correlations: for example, as a thought experiment, imagine trait A and trait B both depend on feature 1, then obviously perception of A and B will have significant correlation. In terms of the LTAs, this means that the LTA for A and LTA for B point in non-orthogonal directions. If LTA for A and LTA for B are separated by an angle less than 90 deg, then we expect them to induce judgments that are positively correlated; otherwise we expect them to induce negatively correlated judgments.

It is perhaps less obvious how data correlation can also induce judgment correlation: as another thought experiment, imagine trait A depends on feature 1, and trait B depends on an orthogonal feature 2. If, for some reason, the joint distribution of feature 1 and feature 2 is positively (negatively) correlated in the natural population of human faces, then there would be a positive (negative) correlation between trait A and B even though they depend on completely unrelated facial features. In terms of LTAs, the data correlation would be apparent in the 2-dimensional subspace spanned by the two LTAs.

Formally, let X *ϵ* ℝ^*D*×*N*^, where *N* is the number of faces, the correlation between the two perceived social traits ŷ_*i*_ and ŷ_*j*_ for the *i*-th and *j*-th traits is given by:

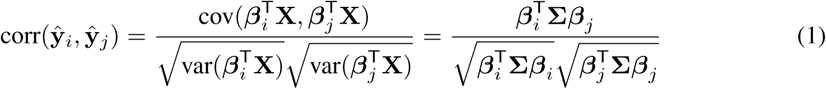

where ∑ is the covariance matrix of X. This ∑ is a diagonal matrix with the variances along each PC on the diagonal. The correlation between ŷ_*i*_ and ŷ_*j*_ do not only depend on the angles between 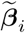 and 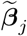, but also the variance of the different dimensions of the PCs. We quantity the “shared feature” component of judgment correlation by the dot product of two LTAs (cosine of the angle between them), which gives us the expected perceptual correlation if the data were isotropically distributed (no data correlation). To quantify the “data correlation” component of judgment correlation, we consider only the 2-dimensional subspace spanned by the two LTAs. The correlation depends sensitively on the coordinate system used. We define the coordinate system, first using 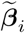 as the first basis vector, and 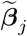 orthogonalized relative to 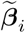 (denoted as 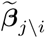) as the second basis vector, and then project all the data (AAM features of face images) into this coordinate system and compute the correlation coefficient. We also use 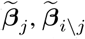 as the coordinate system, and compute the correlation coefficient. We then average the two correlation coefficients to estimate the expected judgment correlation due to data correlation.

Figure 4 shows measured judgment correlation between attractiveness and each of dominance, trustworthiness, femininity, masculinity, babyfacedness, and age (dataset D-2), along with model-predicted correlation, correlation due to shared facial features, and correlation due to data correlation. Note that here, we learned the attractiveness LTA separately for male (A) and female (B) faces. Note that many of these traits show non-zero data correlation with attractiveness, making data correlation an important contributor to judgment correlation. In particular, the relationship between attractiveness and babyfacedness is very interesting. While male and female faces show similar judgment correlation with attractiveness in both actual ratings and model predictions, it turns out that the causes are very different between male and female faces. For male faces, there is almost no shared facial features, and the judgment correlation is primarily driven by data correlation. For female faces, shared facial features contribute significantly to judgment correlation, but there is a strong negative data correlation that counteracts against the first factor, resulting in an overall a relatively weak judgment correlation. Figure 4C;D illustrate these points using the data projected to the 2D subspace spanned by the two LTAs.

**Figure 4:**
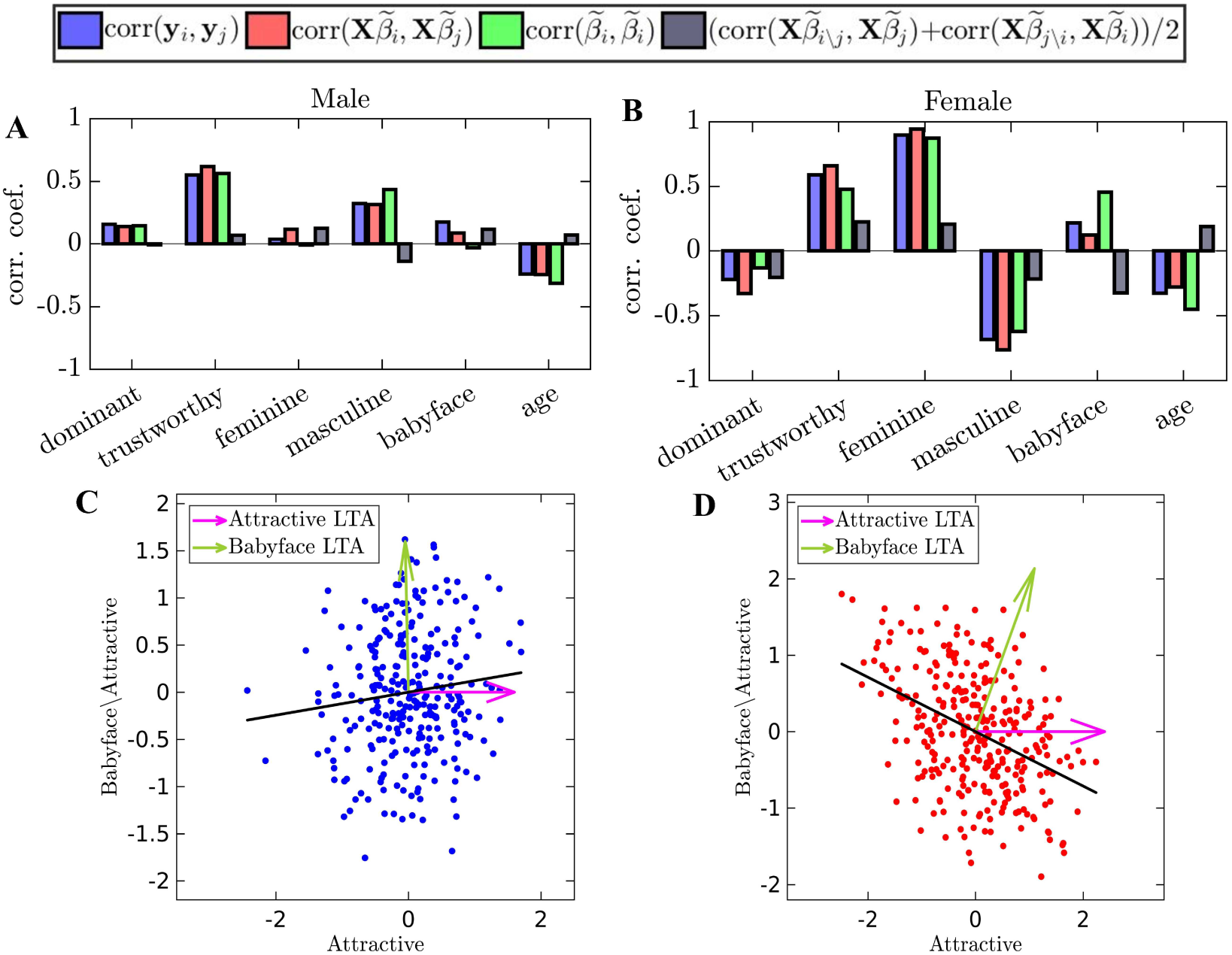
Decomposition of correlations between the perception of attractiveness and babyfacedness for male (A) and female (B) faces in D-2. (C) Projection of faces into the 2D subspace spanned by babyfacedness LTA (green) and attractive LTA (magenta) for male faces. (D) Same as (C) for female faces. Black line: regression line.

Another interesting trait is femininity. We see that while femininity is a very strong predictor of attractiveness in female faces, femininity is barely correlated with attractiveness in male faces, and what little positive correlation there is almost entirely due to data correlation. Indeed, masculinity is strongly correlated with attractiveness in male faces, driven entirely by shared facial features (the data correlation is negative).

## 4 Discussion

In this paper, we showed that our model achieves state-of-the-art performance (comparable to the best CNN) in predicting human social perception on faces, but with a representation that is general purpose, statistical, generative, and transparent. Compared to previous models, it is also more systematic and interpretable. We also presented a number of techniques that allow us to greater insight into the specific combinations of facial features that drive the percept of any particular trait, as well as the relationship among different traits.

Despite our success, there are several necessary directions of future research. The first is that our model needs to be extended to capture nonlinear elements of face perception, such as the classical beauty-in-averageness effect and its contextual modulations [32]. Much work also remains to be done to interpret LTAs in terms of more basic descriptors. The geometric feature analysis was revealing, but much variance remained unexplained, which could be due to nonlinear contributions of these features, other un-measured geometric features, or non-geometric features. Human face processing also appears to be culturally dependent [33] and experience-dependent [34], elements that the model would need to capture.

